# Use of a highly specific kinase inhibitor for rapid, simple and precise synchronization of *Plasmodium falciparum* and *Plasmodium knowlesi* asexual stage parasites

**DOI:** 10.1101/2020.04.24.059493

**Authors:** Margarida Ressurreição, James A. Thomas, Stephanie D. Nofal, Christian Flueck, Robert W. Moon, David A. Baker, Christiaan van Ooij

**Author notes:** Signalling in Apicomplexan Parasites Laboratory, The Francis Crick Institute, London, UK.

## Abstract

During the course of the asexual erythrocytic stage of development, *Plasmodium* spp. parasites undergo a series of morphological changes and induce alterations in the host cell. At the end of this stage, the parasites exit the host cell, after which the progeny invade a new host cell. These processes are rapid and occur in a time-dependent manner. Of particular importance, egress and invasion of erythrocytes by the parasite are difficult to capture in an unsynchronized culture, or even a culture that has been synchronized to within hours. Therefore, precise synchronization of parasite cultures is of paramount importance for the investigation of these processes. Here we describe a method for synchronizing *Plasmodium falciparum* and *Plasmodium knowlesi* asexual blood stage parasites with ML10, a highly specific inhibitor of the cGMP-dependent protein kinase (PKG) that arrests parasite growth approximately 15 minutes prior to egress. This inhibitor allows parasite cultures to be synchronized to within minutes, with a simple wash step. Furthermore, we show that parasites remain viable for several hours after becoming arrested by the compound and that ML10 has advantages over the previously used PKG inhibitor Compound 2. Here, we demonstrate that ML10 is an invaluable tool for the study of *Plasmodium spp*. asexual blood stage biology and for the routine synchronization of *P. falciparum* and *P. knowlesi* cultures.

## INTRODUCTION

The asexual erythrocytic stage of the malaria parasite life cycle is marked by time-specific changes in parasite morphology and modifications to the host cell, driven by a just-in-time gene expression programme in the parasite [1-3]. Currently *P. falciparum, P. knowlesi* and *P. cynomolgi* are the only *Plasmodium spp*. that can be cultured *in vitro* [4-7], and to study these time-dependent changes in culture, the synchrony of the parasites needs to be artificially maintained. Whilst parasites remain synchronous *in vivo* [8-10], the absence of host interactions that regulate the synchrony of parasite development *in vitro* [8, 11] cause the parasites in the culture to become asynchronous – driven by small differences in the length of the replication cycle between individual parasites. This hinders the study of stage-specific processes, including host cell modification, formation of internal organelles, initiation of sexual differentiation, egress and invasion. Some of these processes are very rapid; egress, starting with rounding of the parasitophorous vacuole, followed by lysis of the parasitophorous vacuole membrane and finishing with the lysis of the erythrocyte plasma membrane, takes approximately 10 minutes [12-14], whereas merozoite invasion of fresh erythrocytes takes around 30 seconds for *P. falciparum* [15]. Hence, capturing these events and studying them biochemically requires parasites in culture to be tightly synchronized.

To overcome the problem of asynchrony in *in vitro* cultures, multiple methods of synchronization have been developed. Some of these methods interfere with the DNA synthesis pathway, arresting the parasites at the G1/S transition of the cell cycle. For example, the reversible DNA synthesis inhibitor aphidicolin has been used to arrest parasites prior to entry into S phase [16]. Similarly, treatment with DL-α-difluoromethylornithine depletes polyamines, inducing arrest at the G1/S transition [17, 18]. Addition of polyamines overcomes this arrest, allowing the parasites to enter the G1 phase of the cell cycle in a highly synchronized manner. This method has been used successfully to investigate the cell cycle in *P. falciparum* [18]. Other methods take advantage of the increased permeability of the plasma membrane of erythrocytes infected with mature stage parasites. Treatment of a culture with osmolytes such as D-sorbitol and D-mannitol lyses erythrocytes infected with trophozoite- and schizont-stage parasites, without affecting erythrocytes infected with ring-stage parasites [19]. This has the disadvantage that the developmental stage of the remaining parasites can still vary widely, as even after 18 hours post-invasion (in the case of *P. falciparum*) lysis with these osmolytes is inefficient. Infected erythrocytes containing late-stage *P. knowlesi* parasites can be selectively lysed using guanidine hydrochloride, as they are less sensitive to sorbitol treatments [20]. Similarly, treatment with the pore-forming toxin Streptolysin O has been described as a method to selectively remove erythrocytes infected with late-stage parasites [21]. Furthermore, a microfluidics device has been developed for the enrichment of ring-stage parasites [22]. Other more commonly used methods take advantage of the decreased density of infected erythrocytes containing more mature stages, such as density centrifugation using a Percoll or Nycodenz cushion, to separate erythrocytes infected with later stage parasites from erythrocytes infected with ring-stage parasites and uninfected erythrocytes [23-26]. These schizonts can then be allowed to invade fresh erythrocytes for a limited time followed by removal of the remaining late-stage parasites with a second cushion flotation and/or a treatment with D-sorbitol. This enables establishment of culture with a known synchronicity [27, 28]. However, to obtain a useful parasitemia, an invasion period of several hours is usually required, which is often too long for the precise investigation of rapid processes such as egress and invasion.

Other methods take advantage of the paramagnetic properties of the hemozoin crystal that is formed during the breakdown of hemoglobin to separate erythrocytes infected with late-stage parasites from uninfected erythrocytes and erythrocytes infected with younger parasites [29-31]. Parasites that have started producing hemozoin will adhere to the matrix of a magnetic-activated cell sorting (MACS) column while the younger parasites and uninfected erythrocytes pass through. This method yields a very high percentage of infected cells, but as the formation of the paramagnetic hemozoin crystal starts relatively early in the erythrocytic cycle, the isolated parasites can vary in age by many hours.

To provide higher degrees of synchrony, various other methods have been developed. For instance, by overlaying schizont-infected erythrocytes immobilized using concanavalin A with uninfected erythrocytes, newly infected erythrocytes can be harvested from the supernatant [32], thereby allowing parasites to be synchronized to within a thirty-minute window.

The method of synchronization with the best temporal resolution is the filtration method. Here, parasite egress is blocked with the irreversible cysteine protease inhibitor E64, followed by mechanical lysis of infected erythrocytes by filtration. In the case of *P. falciparum*, parasites are passed through a 1.5 μm filter, whereas for *P. knowlesi* double filtration through a 3 μm filter followed by 2 μm filter is applied, resulting in a nearly instantaneous release of infectious merozoites [33-35]. This method provides a resolution of invasion of seconds to minutes and has provided important insights [36]. However, the proportion of viable merozoites produced by this method is highly variable and can be quite low. So whilst well-suited for comparing the effect of treatments on a single parasite strain (e.g. drug treatments), comparisons between different parasite strains are challenging (such as comparing mutant and wild-type parasites). This precludes its use for larger-scale investigations and assays and would be cumbersome to apply to routine synchronization of parasite cultures.

To overcome these drawbacks, chemical inhibition of cyclic GMP (cGMP)-dependent protein kinase (PKG) has been used to synchronize parasites. Activation of PKG initiates a signaling cascade that promotes the lysis of the parasitophorous vacuole membrane and the erythrocyte membrane; inhibition of this kinase arrests the parasites at a step immediately prior to the initiation of egress [37-40]. Subsequent removal of reversible PKG inhibitors leads to rapid release of merozoites. One such inhibitor, an imidazopyridine referred to as Compound 2 (4-[7-[(dimethylamino)methyl]-2-(4-fluorophenyl)imidazo[1,2-α]pyridine-3-yl]pyrimidin-2-amine) [41], has been applied to the study of egress and invasion. The timed release of a synchronized culture of parasites allows these rapid events to be visualized which are normally extremely difficult to observe, even in a population where the window of synchronization is hours [13, 14, 38, 42-45]. However, a detailed description of the use of PKG inhibitors has not been published. Recently a highly specific and potent derivative of Compound 2, referred to as ML10, with a single digit nM EC_50_ and an approximately 75-fold higher specificity has been described [46]. To demonstrate that ML10 is a valuable tool for the synchronization of *Plasmodium* spp. parasites, we determined a number of important parameters for the use of this compound.

## RESULTS

### Working concentration of ML10 for *P. falciparum* and *P. knowlesi*

To develop ML10 as a reagent for the synchronization of malaria parasites, we first derived an optimal working concentration. The EC_50_ of ML10 on the *P. falciparum* strain 3D7 was determined using a standard 72-hour SYBR Green growth inhibition assay, starting with young trophozoite-stage parasites [47]. With this assay, an EC_50_ of 4.2 nM (±SD 2.7 nM) was obtained (Figure 1A, Table 1, Supp. Figure 1A), very similar to the EC_50_ of 2.1 nM (±SD 0.2 nM) determined previously using the hypoxanthine incorporation assay [46]. The EC_50_ of Compound 2 in the same SYBR Green 72-hour growth inhibition assay was 100 nM (±SD 35 nM), also similar to previously published values using the hypoxanthine incorporation assay (Figure 1B, Supp. Figure 1B) [46]. To verify the specificity of the inhibitors for PKG in the SYBR Green assay, we determined in parallel the EC_50_ for a 3D7-derived transgenic line that harbors a gatekeeper mutation (T618Q) in the PKG sequence [40, 46]. In this mutant, access to the ATP-binding pocket is blocked, preventing binding of ATP-competitive PKG inhibitors that utilise the gatekeeper pocket, including ML10, Compound 2 and Compound 1, however, this mutation has no detectable effect on the endogenous activity of PKG or parasite growth [41, 46, 48]. The EC_50_ values of ML10 and Compound 2 in this strain were several orders of magnitude higher than that of 3D7 (Figures 1C and 1D), confirming that ML10 is indeed a highly specific inhibitor of PKG, as previously described [46]. As the results using the 72-hour SYBR Green growth inhibition assay closely replicated the data previously obtained with the hypoxanthine incorporation assay, it was deemed suitable for further investigation of ML10. Based on these results, we set the working concentration of ML10 for subsequent experiments at 25 nM, a concentration approximately six-fold higher than the EC_50_ for wild type parasites, but that had no effect on the gatekeeper mutant parasites. From this we conclude that at this concentration, ML10 does not affect any essential target other than PKG.

**Table 1.** EC50 of ML10 for different *Plasmodium* species grown in RPMI-1640 containing various medium supplements.

| <u>Species</u> | <u>Strain</u> | <u>Medium supplement</u> | <u>EC50 ± SD</u> | <u>Working concentration</u> |
| --- | --- | --- | --- | --- |
| *Plasmodium falciparum* | 3D7 | AlbuMax | 4.2 ± 2.7 nM | 25 nM |
| *Plasmodium falciparum* | 3D7 | AlbuMax, 10% HoS | 17.9 ± 3.3 nM | 50 nM |
| *Plasmodium falciparum* | HB3 | AlbuMax | 4.9 ± 2.9 nM | 25 nM |
| *Plasmodium falciparum* | Dd2 | AlbuMax | 5.1 ± 1.5 nM | 25 nM |
| *Plasmodium falciparum* | NF54 | AlbuMax | 6.9 ± 2.0 nM | 25 nM |
| *Plasmodium knowlesi* | A1-H.1 | AlbuMax, 10% HoS | 43.8 ± 5.6 nM | 150 nM |

**Figure 1.**
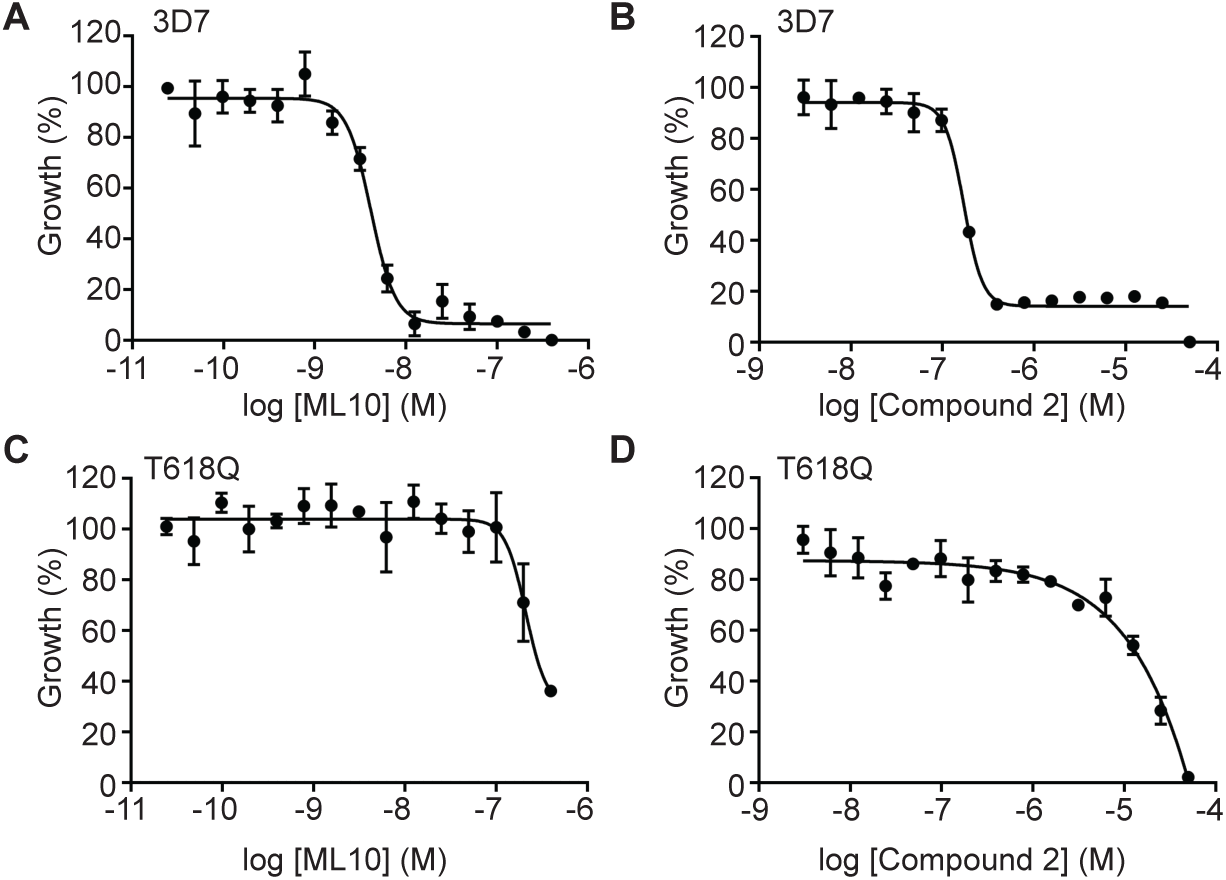
Dose-response curves of ML10 and Compound 2 for *P. falciparum* 3D7 and T618Q parasites. *P. falciparum* 3D7 parasites (**A** and **B**) and T618Q parasites that harbor a gatekeeper mutation in the gene encoding PKG (**C** and **D**) were cultured in the presence of decreasing concentrations of ML10 (**A** and **C**) or Compound 2 (**B** and **D**). Parasite growth was determined using a 72-hour SYBR Green assay and the EC_50_ value for the compounds was calculated for each strain from the resulting dose-response curve. Shown are representative graphs from one of five replicates. For EC_50_ values of all five experiments, see Supp. Figure 1; for the average EC_50_ values, see Table 1. For both compounds, the difference between the EC_50_ for 3D7 and the T618Q parasites was statistically significant by an unpaired t-test, in both cases, *p*≤0.001.

To investigate whether the 25 nM working concentration used for 3D7 would be widely applicable to other *P. falciparum* isolates, we determined the EC_50_ of ML10 in three additional *P. falciparum* laboratory isolates from different parts of the world and with different drug resistance phenotypes [49]: Dd2 (originally from Southeast Asia, has increased resistance to chloroquine, cycloguanil and pyrimethamine), HB3 (originally from Honduras, has increased resistance to cycloguanil and pyrimethamine) and NF54 (isolated from a case of ‘airport malaria’; this strain has similarities to African parasite strains [50]) from which 3D7 is derived. The EC_50_ values obtained for these strains were 5.14 nM (±SD 1.5 nM) for Dd2, 4.86 nM (±SD 2.9 nM) for HB3 and 6.87 nM (±SD 2.0 nM) for NF54 (Figures 2A, 2B and 2C, Table 1, Supp. Figure 1A). The EC_50_ of Compound 2 in these strains was determined to be 150 nM (±SD 46 nM) for Dd2, 104 nM (±SD 36 nM) for HB3 and 130 nM (±SD 12 nM) for NF54 (Figures 2D, 2E and 2F, Table 1, Supp. Figure 1B). Hence, ML10 and Compound 2 affect these distinct *P. falciparum* laboratory isolates similarly and therefore the 25 nM working concentration is likely to be effective for many, if not all, *P. falciparum* isolates.

**Figure 2.**
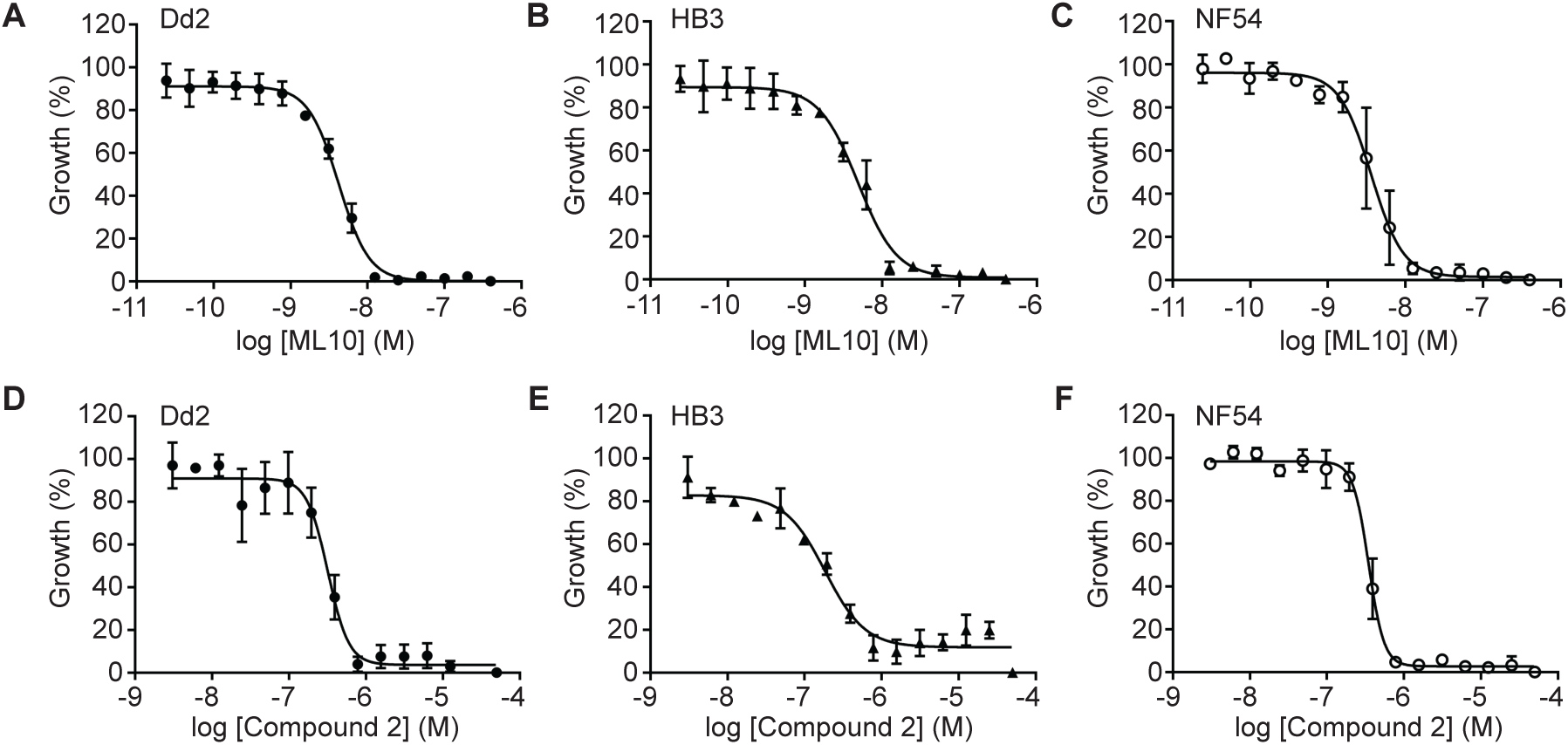
Dose-response curves of *P. falciparum* strains Dd2, HB3 and NF54 cultured in the presence of ML10 or Compound 2. *P. falciparum* strains Dd2, HB3 and NF54 were cultured in the presence of decreasing concentrations of ML10 (**A-C**) or Compound 2 (**D-F**) for 72 hours and growth was determined by staining with SYBR Green. The resulting dose-response curves were used to calculate EC_50_ values for the compounds. Shown are representative dose-response curves converted to percentage from one of three independent experiments. For EC_50_ values of all experiments, see Supp. Figure 1. Difference in EC_50_ values between the strains were determined not to be significant using a Kruskal-Wallis one-way analysis of variance.

Next, we established the EC_50_ of ML10 for *P. knowlesi*, the only other human malaria species that can be cultured *in vitro* in human red blood cells [5, 6], using the same SYBR Green growth assay. Surprisingly, the EC_50_ of ML10 for this species was determined to be 43.8 nM (±SD 5.6 nM) (Figure 3A), over ten-fold higher than for *P. falciparum*. Whereas *P. falciparum* is grown in medium containing only 0.5% AlbuMax, *P. knowlesi* is cultured *in vitro* in a high-serum medium containing both 10% horse serum and 0.5% AlbuMax, and this additional protein binding content is known to increase EC_50_ values owing to the binding of compounds to serum components [51]. To determine if this was the case for ML10, we measured the EC_50_ of ML10 for *P. falciparum* 3D7 grown in the horse serum-containing medium used to culture *P. knowlesi*. This revealed that the addition of serum increases the EC_50_ of ML10 in *P. falciparum* 3D7 parasites to 17 nM (Figures 3B, Table 1, Supp. Figure 1B), approximately four-fold higher than when grown without serum – indicating the serum content of the medium accounted for the majority of the differential susceptibility. We therefore tested whether a concentration of 150 nM ML10 could arrest a *P. knowlesi* culture without affecting the viability of the parasites. Treatment of a *P. knowlesi* culture did indeed arrest parasites at the end of the cycle (Figure 3C). When ML10 was removed when ring-stage parasites were detected in the DMSO-treated control samples (approximately two hours after the addition of ML10), the parasites were able to egress and invade new erythrocytes, demonstrating that they remained viable after ML10 treatment (Figure 3C). As *P. falciparum* parasites are also often cultured in the presence of human serum, we further tested whether 25 nM, 50 nM or 75 nM ML10 would be an effective concentration to arrest *P. falciparum* grown in the presence of human serum. We found that when cultured in either the presence of 10% human serum (Figure 3D) or 5% human serum in addition to 0.5% AlbuMax (Figure 3F), 25 nM ML10 did not effectively block progression of the parasites, whereas 50 nM and 75 nM ML10 did. Hence, using a concentration of 50 nM ML10 in cultures containing human serum is sufficient to arrest parasites (Table 1).

**Figure 3.**
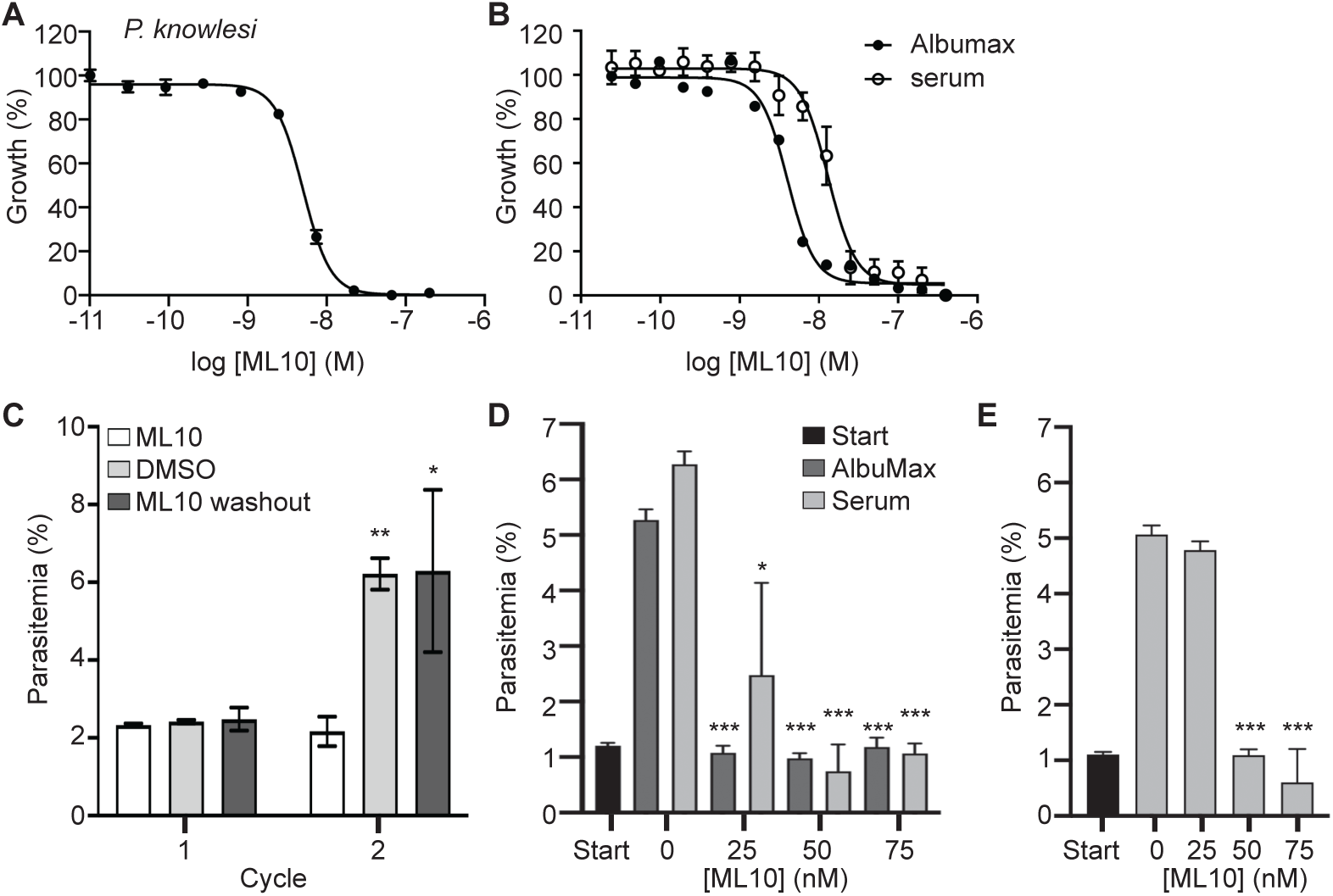
Dose-response curve of *P. knowlesi* cultured in the presence of ML10 and effect of serum on EC_50_ of ML10. Dose-response curves derived from parasite growth inhibition assays obtained with *P. knowlesi* (**A**) and *P. falciparum* strain 3D7 cultured in RPMI-1640 with or without supplementation with 10% horse serum (**B**). These results were used to calculate the EC_50_ of ML10 on parasites during the erythrocytic stage. (**C**) Effect of 150 nM ML10 on the growth of *P. knowlesi* egress. Late stage synchronized parasites were incubated in the presence of 150 nM ML10 (bars on left-hand and right-hand side) or DMSO (bars in middle). When ring-stage parasites were detected in the untreated sample (approximately 2 h after addition of ML10), the parasites were washed to remove the ML10 or DMSO. Parasites were subsequently incubated in the presence of DMSO or ML10. Note that 150 mM ML10 completely blocks progression to the next cycle, whereas removal of the compound after two hours of incubation has no effect on the parasitemia in the next cycle. Results of statistical comparison of the ML10-treated sample with the DMSO and the wash-out sample in each cycle are indicated: no symbol: no significant difference; *: *p*≤0.05; **: *p*≤0.01. (**D, E**) To identify an effective working concentration of ML10 for *P. falciparum* in the presence of serum, parasite cultures were cultured in the presence of 0.5% AlbuMax alone or AlbuMax and 5% serum (**E**) or 10% serum (without additional AlbuMax) (**F**) and ML10 at concentrations ranging from 0-75 nM for one cycle. Start (black bars) denotes the parasitemia of the culture in the first cycle, prior to the addition of ML10. Error bars denote the standard deviation of three replicates. Results of statistical comparison of 0 nM sample with the samples treated with ML10 is indicated: *: *p*≤0.05; \*\*\**p*≤0.001.

### Schizont rupture arrested with ML10

As ML10 reversibly blocks the intra-erythrocytic parasite at a stage immediately prior to the onset of egress, it is an ideal reagent to synchronize egress and the subsequent invasion of merozoites to obtain highly synchronized cultures. However, synchronization of an asynchronous parasite culture requires that the most mature parasites be held in an arrested state for an extended time until the younger parasites reach the same arrested state. To determine the length of time that arrested parasites remain viable, we tightly synchronized a 3D7 culture with two sequential flotations on a Percoll cushion. Approximately 45 hours later, when rings were detected in the culture, schizonts were isolated and mixed with uninfected erythrocytes to give a final parasitemia of ∼1%. This culture was split into three flasks, to which either DMSO, 25 nM ML10 or 1.5 μM Compound 2 was added. These cultures were then incubated at 37°C for 1-12 h. At the end of the incubation period, the parasites were pelleted and resuspended in fresh medium without compound, allowing viable parasites to egress and invade fresh erythrocytes. The resulting parasitemia serves as a measure of the survival of the schizonts during the incubation in the presence of ML10 and Compound 2. This revealed that incubation in the presence of ML10 for up to three hours did not affect the viability of the parasites and that the effect of ML10 on parasite viability after four hours was generally small (Figure 4A). However, a noticeable decline in viability was detected after six hours, with further declines as the length of the incubation increased. As the viability of the parasites decreased, they took on a bloated appearance in Giemsa-stained blood films, in contrast to the healthy appearance of parasites seen in the samples treated for shorter periods (Figure 4B). Interestingly, the survival of parasites incubated in the presence of Compound 2 was consistently lower (Figure 4A). Nonetheless, the viability of parasites incubated in the presence of Compound 2 showed a similar decrease in viability over time as those incubated with ML10: decreasing after four hours in the presence of the compound, with larger decreases starting after six hours on incubation. The higher level of survival of parasites in the presence of ML10 as compared to Compound 2 indicates that this compound may be a superior choice for the synchronization of parasites.

**Figure 4.**
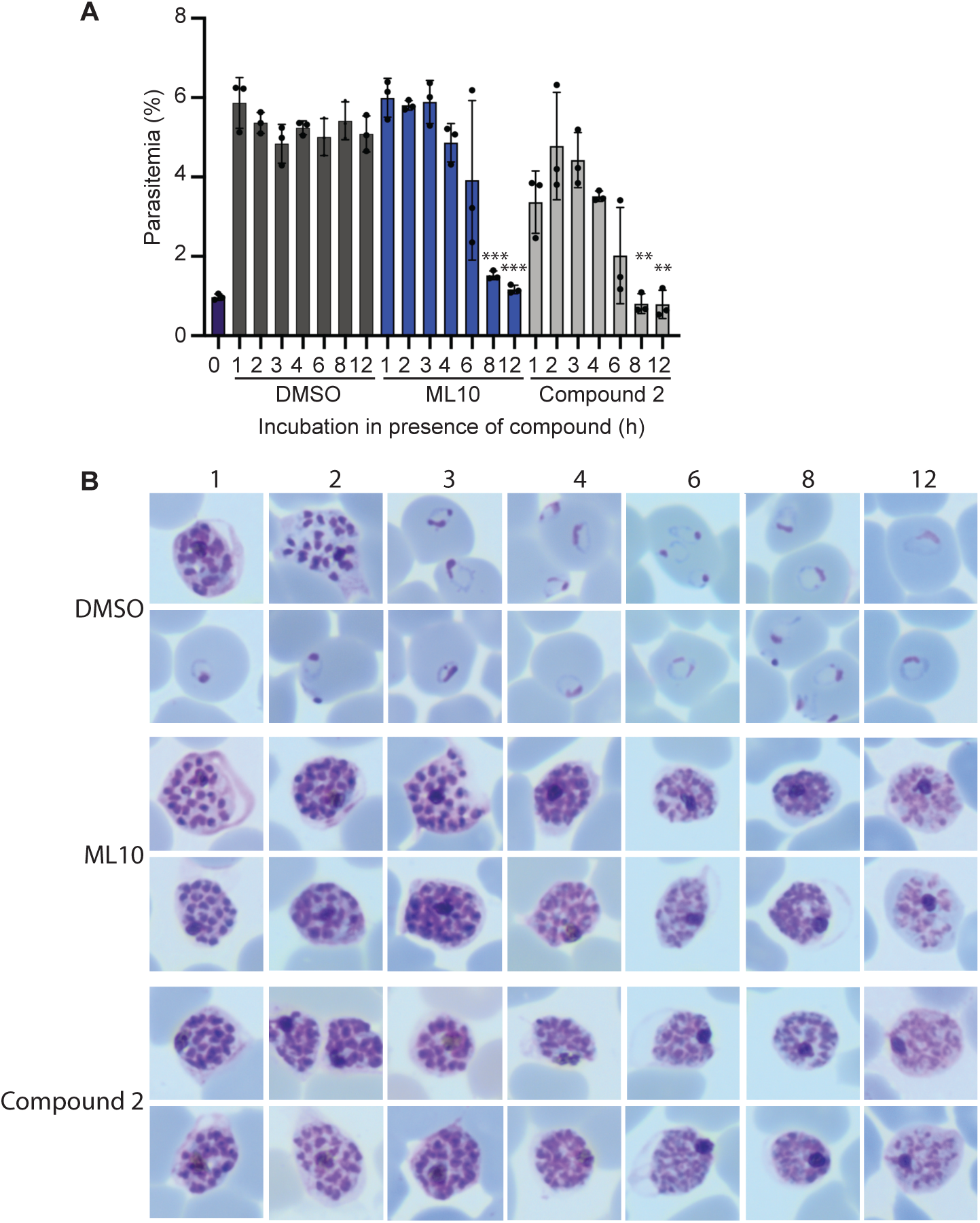
Impact of the length of exposure of arrested mature schizonts to ML10 or Compound 2 on parasite viability. (**A**) Cultures containing mature schizonts were maintained for 0-12 hours in either DMSO, 25 nM ML10 or 1.5 µM Compound 2 and subsequently washed and allowed to egress and invade fresh red blood cells. The resulting parasitemia was determined by flow cytometry and reflects the survival of the parasites in the compound. Shown is a representative experiment of five independent experiments; each experiment consisted of three technical replicates. The statistical significance of the difference between the parasitemia of the culture treated for one hour and the subsequent samples was determined using an unpaired t-test; ** = *p*≤0.01, *** = *p*≤0.001, whereas no symbol indicates no statistical significance. (**B**) Giemsa-stained parasite smears were produced immediately prior to removal of the compound to ascertain the effect of incubation in the presence of the compound on the morphology of the parasites. Numbers above the images denote the length of time (in hours) the parasites had been exposed to the compound.

### Timing of egress after removal of ML10 and synchronicity of resulting cultures

Induction of PKG activity initiates a cascade of downstream events that ultimately results in egress [37, 38, 40], causing a delay between the initiation of PKG activity after the removal of ML10 and egress. Video microscopy was used to monitor egress of closely synchronized late schizonts arrested with ML10 or Compound 2 and following the removal of the compounds. When ML10 was tested, on average the first parasites began to egress 17 mins 18 s (±SD 3 min 38 s) after ML10 washout and many of the parasites that egressed in the course of the imaging were detected within an 11 minute window (±SD 4 min 16 s) (Supp. video 1 and Supp. Figure 2). When Compound 2 was tested in this assay, release of parasites was detected in a similar timeframe, with the first parasites egressing on average 16 mins 48 s (±SD 5 min 7 s) after compound removal and the remaining parasites that egressed in the course of the imaging were released within a 13 minute and 50 s window (±SD 2 min 45 s SD) (Supp. video 2 and Supp. Figure 2). This is comparable to previously reported timings of parasite release following the removal of Compound 2 [14, 38, 42].

The results of the video microscopy indicated that the burst of merozoite release was very synchronous, although not all parasites egressed during the imaging period. This may reflect that these parasites had not yet matured to the stage where they were blocked by either compound at the time it was removed or that a fraction.

To determine the synchronicity of invasion after removal of ML10, we measured the appearance of ring-stage parasites in the culture by flow cytometry at 10-min intervals after the removal of ML10 or Compound 2. Most egressed parasites were detected 20 min after the removal of the compound, with only very small increases afterwards, indicating that egress of parasites after removal of the compounds occurs in a narrow time window and results in a highly synchronous culture (Figures 5A and 5B). Whereas it was difficult to distinguish attached from invaded merozoites at the 20-minute time point, at the 30-minute time point, the parasites had progressed to the typical ring-shaped morphology of very young parasites, indicating that they had entered the erythrocyte. Interestingly, treatment with Compound 2 resulted in fewer ring-stage parasites compared to treatment with ML10, similar to what was observed when parasites were incubated in ML10 or Compound 2 for extended periods of time (Figure 4), again potentially indicating that there are off-target effects of Compound 2, although the timing of the appearance of new rings and the synchrony were very similar between the two compounds.

**Figure 5.**
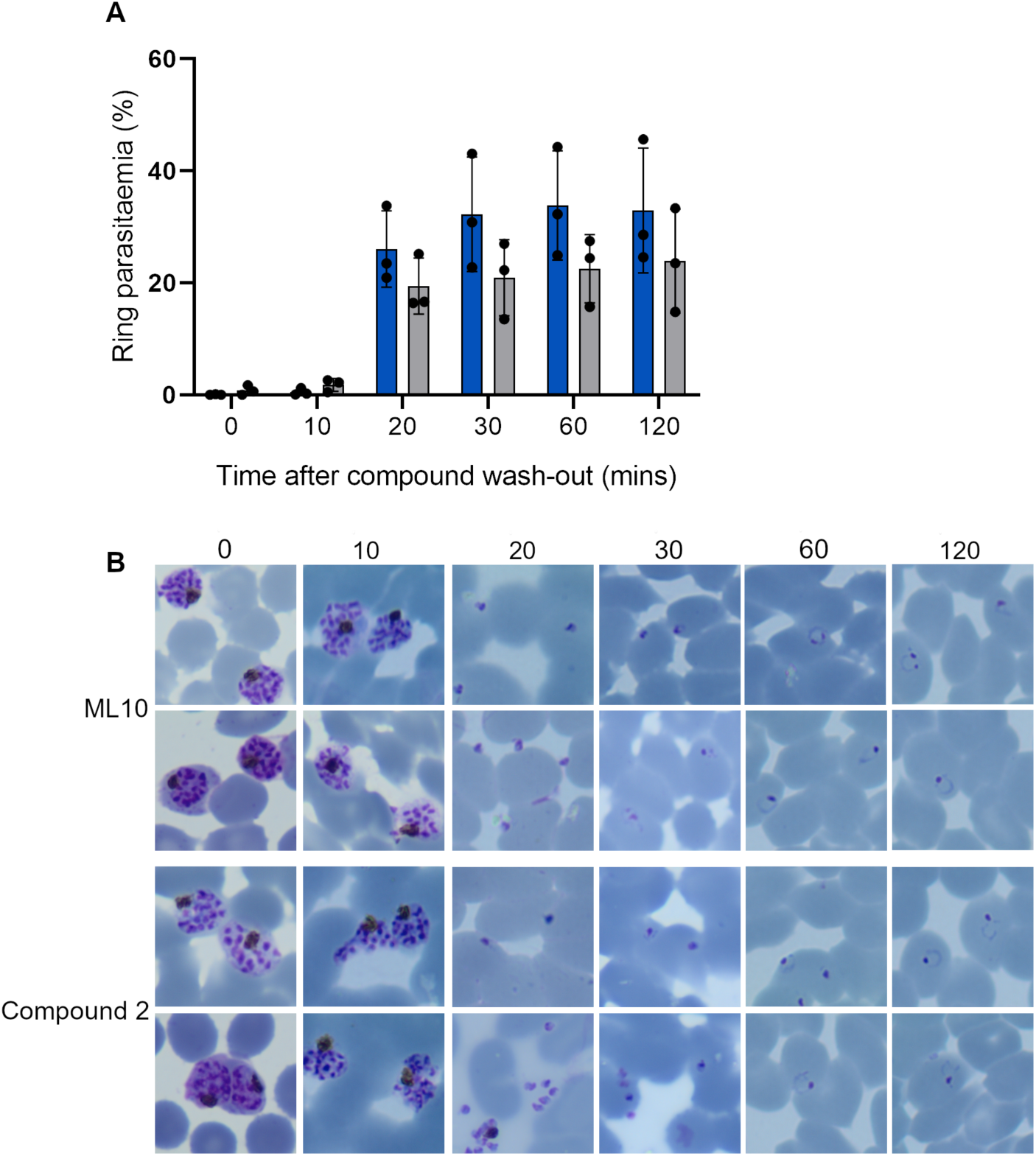
Synchronicity of invasion after removal of inhibition by ML10 or Compound 2. (**A**) Parasites that had been arrested with either ML10 (blue bars) or Compound 2 (grey bars) for up to three hours were washed to allow egress and invasion of new erythrocytes. Samples were removed at 10-minute intervals and the percentage of host cells infected with ring forms, which represent parasites that have egressed and invaded new erythrocytes, was determined by cytometry (see Supp. Figure 3 for gating strategy). Error bars represent standard deviation from 3 independent experiments each conducted in duplicate, no statistically significant differences were detected between compounds at each time point using a pairwise individual t-test. (**B)** Giemsa smears made at the time point indicated at the top confirmed the presence of ring-stage parasites in the samples.

### Development of parasites through the erythrocytic cycle in the presence of ML10

All of the above experiments were performed by adding ML10 to mature schizonts, either in culture or purified by flotation on a Percoll cushion. However, for the continued maintenance of a synchronous culture, it would be beneficial if the compound could be added at any stage of the cycle. To test if ML10 affects any other part of the intraerythrocytic cycle, we added either DMSO or ML10 to a culture of ring-stage parasites that had invaded within the previous two hours. We subsequently monitored the development of the parasites by Giemsa staining and by measuring the increase in DNA content of the parasites by flow cytometry. This revealed no obvious difference between the DMSO-treated and ML10-treated parasites in their progression through the intraerythrocytic cycle (Figures 6A). To examine if viable merozoites were produced by parasites that had developed for the entire cycle in the presence of ML10, DMSO and ML10 were removed from the cultures three hours after ring-stage parasites were first detected in the DMSO-treated culture. At this time, the parasites in the ML10-treated culture had been blocked for up to three hours by the compound and hence had completed the entire intra-erythrocytic cycle in the presence of the compound. When the parasitemia of the cultures in the next cycle was determined, no significant difference was detected, indicating that ML10 does not affect the growth of the parasites and the production of viable merozoites even when the compound is present throughout the entire intra-erythrocytic cycle (Figure 6B).

**Figure 6.**
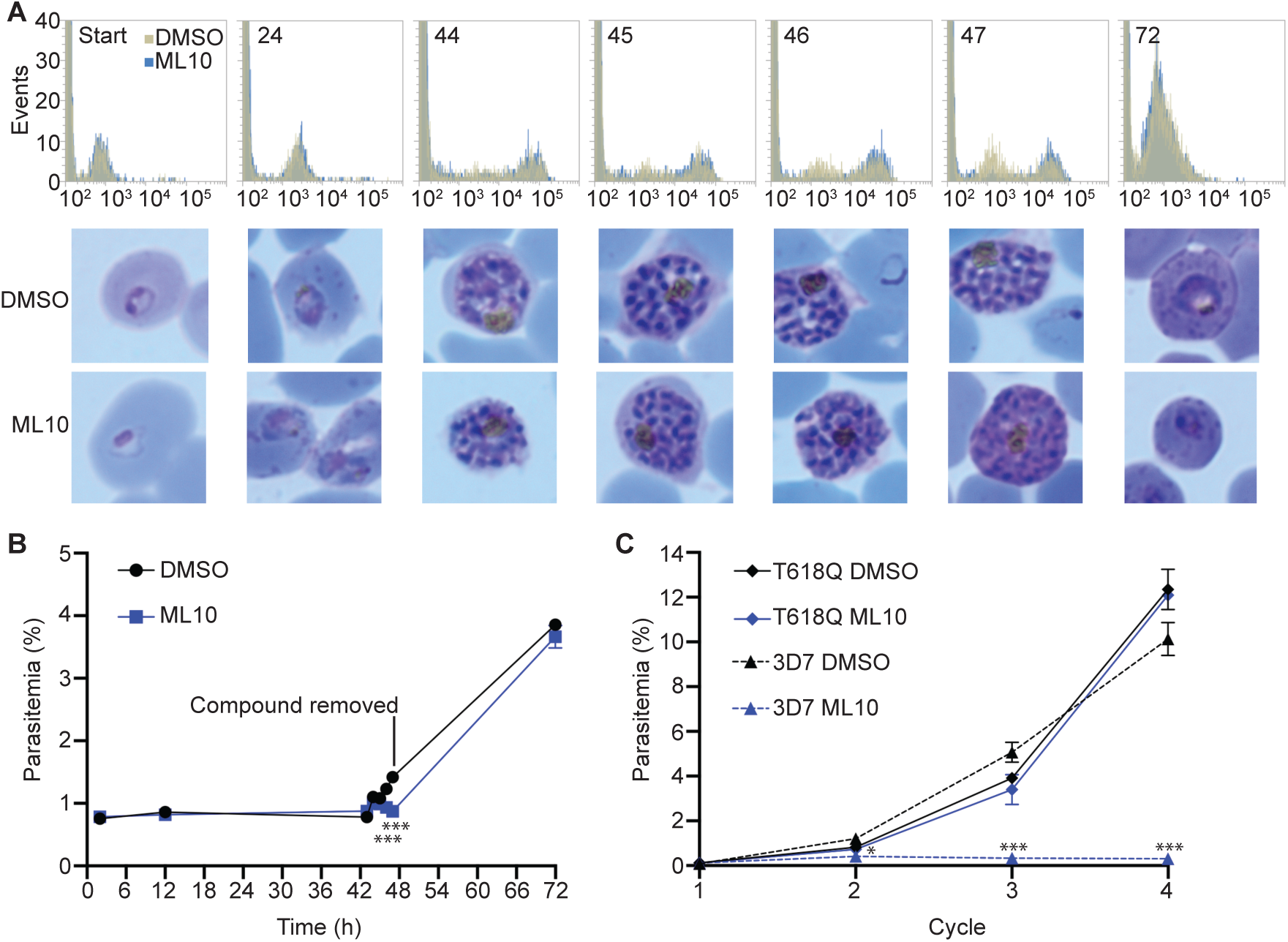
Growth of *P. falciparum* in the presence of ML10. (**A**) Cultures of *P. falciparum* strain 3D7 consisting of rings no older than 2 hours were treated with 25 nM ML10 or an equivalent amount of DMSO and parasite development was monitored throughout the erythrocytic cycle for the synthesis of DNA, as measured by SYBR Green I staining and flow cytometry (top panels; gold represents the DMSO-treated sample, while blue represents the ML-10 treated sample. The number in each panel indicates hours after onset of invasion, while the x-axis shows the nucleic acid content as measured with SYBR Green I staining) and imagining of Giemsa-stained smears (bottom panels). At the 47-hour time point (three hours after ring-stage parasites were first detected in the DMSO-treated samples), DMSO and ML10 were removed. (**B**) The parasitemia of the cultures in (A) was determined at each time point. Note the similar parasitemias of the DMSO-treated and ML10-treated samples in the second cycle, indicating that the presence of ML10 in the first cycle did not affect the development of the parasites. The cultures were set up in triplicate, error bars represent standard deviation at each time point. The statistical significance of the difference between the parasitemia of the DMSO-treated and ML10-treated samples at each time point was calculated using an unpaired t-test; *** indicates *p*≤0.001, whereas no symbol indicates no statistical significance. (**C**) Growth of *P. falciparum* strains 3D7 and T618Q over four cycles. Parasite cultures were diluted to a parasitemia of ∼0.1% and growth was monitored over four cycles. Medium was changed daily after four days. Note the nearly identical growth of the DMSO-treated and ML10-treated T618Q parasites. The cultures were set up in triplicate, error bars represent standard deviation at each time point. The statistical significance of the difference between the parasitemia of the DMSO-treated and ML10-treated samples for each strain at each cycle was calculated using an unpaired t-test; *= *p*≤0.05 and ***=*p*≤0.001, whereas no symbol indicates no statistical significance. The graphs represent one of three biological replicates.

To determine if treatment with ML10 can have subtle effects on the parasites if used over prolonged periods, we tested the effect of continuous treatment with ML10 on the growth of the T618Q strain and 3D7 over several cycles. The T618Q strain was not affected by the presence of ML10, as indicated by the identical growth rate in the presence or absence of ML10, whereas the 3D7 strain grew only in the absence of ML10 (Figure 6C). This further shows that at the concentration used (25 nM) the only essential molecule in the parasite affected by ML10 is PKG.

## DISCUSSION

The data presented here provide the first in-depth analysis of the use of PKG inhibitors as tools to synchronise and study the intra-erythrocytic cycle of *Plasmodium* spp. parasites. We show that the PKG inhibitor ML10, a derivative of the previously used imidazopyridine PKG inhibitor Compound 2 [46], reversibly arrests *P. falciparum* and *P. knowlesi* at a very late schizont stage at low nanomolar concentrations. Whereas we detected no significant difference in the EC_50_ of ML10 among four different *P. falciparum* laboratory isolates, the EC_50_ of ML10 was markedly influenced by the supplement used in the medium; human and horse serum increase the EC_50_ compared with medium containing AlbuMax, likely owing to binding of the compound to serum components. The EC_50_ for *P. knowlesi* was found to be higher than that for *P. falciparum*, even when *P. falciparum* was grown in the presence of serum, indicating a difference in biology that may make it slightly less susceptible to PKG inhibition. This is similar to previous findings, which established that the EC_50_ values of certain anti-malarials are higher for *P. knowlesi* than for *P. falciparum* [51, 52]. Nonetheless, we have been able to successfully calibrate concentrations that are effective in the two species and in several media formulations (Table 1).

Removal of ML10 leads to a highly synchronous burst of egress and subsequent invasion, as a large fraction of the parasites released after the removal of the compound had invaded within a ten-minute window. By preventing further invasion through the addition of compounds known to inhibit invasion, such as heparin [53], a culture could be established in which the parasites are matched in age to within minutes.

In contrast to other methods for synchronizing *Plasmodium* cultures, which can be labour-intensive or require specialized equipment, synchronizing cultures with ML10 is fast and simple. Removal of the compound from an arrested culture with a single wash step leads to egress of the arrested parasites and invasion of erythrocytes within 20 minutes. As parasites arrested by ML10 retain their viability for several hours, a semi-synchronized parasite culture can be treated with ML10 for an extended period to allow a subset of parasites to reach the point in the cycle where they become arrested. Subsequent removal of the compound will then cause a proportion of parasites to undergo egress and invade fresh erythrocytes in a very short time span. Parasites that have not reached the point of ML10 arrest, and hence have not yet egressed, can then be removed on a Percoll cushion or by lysis with 5% D-sorbitol, thereby establishing a highly synchronized culture. As ML10 does not affect any essential parasite proteins other than PKG, synchronicity in the culture can then be maintained simply by adding ML10 at any stage of the asexual cycle and removing the compound when the parasites become arrested prior to the initiation of egress. In our experience, simply replacing the medium in a culture flask with fresh medium is sufficient to lower the concentration of ML10 to below its effective concentration and allow egress and subsequent invasion. Hence, addition of ML10 provides a very simple method for the maintaince of synchronicity of parasite cultures over many cycles.

Synchronized cultures are of great value in the study of many processes in the asexual cycle and the ability to synchronize egress and invasion to within minutes will allow the investigation of these rapid processes on different scales, allowing for a wide range of studies, including large-scale biochemical investigation of proteins involved, such as the proteolysis of MSP1 and AMA1 proteins [54-56]. Furthermore, as it makes timing of egress and invasion very predictable, this method will greatly aid the imaging of egress and invasion by video microscopy. In addition, a culture of such tightly synchronized parasites will be of great benefit for the study of very early ring-stage parasites; parasites at this stage undergo very rapid changes [57, 58] that are difficult to capture on a population level in a culture in age by hours. An additional application of the very tight synchronization is the preparation of parasites for schizont-stage transfection, which requires extremely synchronized parasites [6, 59]. Compound 2 has been used previously to aid synchronization of *P. knowlesi* for transfection [44]. Furthermore, ML10 has recently been used to block the proliferation of asexual stages in a *P. falciparum* culture induced to form gametocytes [60].

Of the PKG inhibitors used for synchronization of parasite cultures, Compound 2 has been the most commonly used [14, 38, 42, 44, 45, 54, 55, 61]. However, the results shown here indicate that ML10 has advantages over Compound 2. Owing to its potency, ML10 can be used at extremely low concentrations (25 nM in the case of *P. falciparum* cultured in AlbuMax, compared to a working concentration of 1 μM in the case of Compound 2), allowing the concentration of the DMSO solvent, which is used to dissolve both ML10 and Compound 2, to remain very low as well; high concentrations of DMSO can have deleterious effects on eukaryotic cells [62, 63]. The low working concentration of ML10 also ensures that stock solutions are not rapidly depleted, allowing routine synchronization of large-scale cultures, and the higher specific activity of ML10 compared to that of Compound 2 additionally reduces the likelihood that any phenotype detected is the result of off-target effect of the compound. Furthermore, we consistently saw a small but noticable decrease in the viability of the parasites treated with Compound 2, even after short incubations with the compound (Figures 4 and 5). This loss of viability was not detected in parasites treated with ML10, indicating that Compound 2 may have additional off-target effects on the parasites or is not washed away as readily as ML10.

In conclusion, we believe that ML10 will be a very valuable tool for the investigation of the asexual erythrocytic stage of the lifecycle of *Plasmodium spp*. The compound is available from LifeArc.

## MATERIAL AND METHODS

### Parasite culture

Unless otherwise stated, all *P. falciparum* experiments were performed with the 3D7 strain and all *P. knowlesi* experiments were performed with the A1-H.1 strain [6]. Both species were maintained in RPMI-1640 medium (Life Technologies) supplemented with additional glucose, 0.5% AlbuMax type II (Gibco), 50 μM hypoxanthine and 2 mM L-glutamine (complete medium) at 2-3% hematocrit at 37°C. In the case of *P. knowlesi*, this medium was further supplemented with 10% (v/v) horse serum (PAN Biotech; [64]). *P. falciparum* was maintained in the presence of 5% CO_2_, whereas *P. knowlesi* was maintained in the presence of 96% N_2_, 1% O_2_, 3% CO_2_. Human erythrocytes were obtained from the National Blood Transfusion Service, UK.

### Measurement of EC_50_ by SYBR Green assay

To assess the ML10 concentration required to block *P. falciparum* and *P. knowlesi* egress and compare this to Compound 2, an *in vitro* 72 h SYBR Green growth inhibition assay was adapted from Smilkstein et al. and van Schalkwyk et al. [47, 51]. A starting culture consisting of synchronous young trophozoite *P. falciparum* or *P. knowlesi* parasites was adjusted to 2% parasitaemia and 2% haematocrit. Starting ML10 and Compound 2 concentrations were 2-fold serially diluted in triplicate in 50 µl complete medium in a 96-well plate to provide a concentration range of 400 nM to 0.024 nM for ML10 and 50 μM to 3 nM for Compound 2. Subsequently, 50 µl of culture was added to each well, to produce a final parasitemia and hematocrit of 1%. The highest DMSO concentration present in the assay was 0.05%. The 96-well culture plates were maintained at 37°C for 72 h in the case of *P. falciparum* and 48 h in the case of *P. knowlesi* and subsequently frozen. For lysis and fluorescence analysis, plates were thawed at 37°C and 100 µl of lysis buffer (20 mM Tris pH 7.5, 5 mM EDTA, 0.008% Saponin and 0.08% Triton X-100 in distilled H_2_O) complemented with SYBR Green I Nucleic Acid Stain at a concentration of 2x (from a 10,000 x stock; Fisher Scientific) was added and incubated in the dark at room temperature for 1 h after thorough mixing. Subsequently, relative fluorescent units were recorded using a Spectramax 3 spectrophotometer with excitation and emission settings at 485 and 535 nm, respectively. The data were obtained in relative fluorescent units and fitted to a dose-response curve (GraphPad Prism 8 software). The EC_50_ (here defined as the drug concentration that gives half the maximal response of egress arrest) was determined by dose-response non-linear regression analysis of the log dose-response curves.

To determine if 150 nM ML10 was an appropriate working concentration for *P. knowlesi*, late stage parasites were purified on a cushion of 55% Nycodenz as described previously [6], washed in RPMI 1640 and allowed to egress and infect fresh erythrocytes in complete *P. knowlesi* medium [6] with agitation for 1 h before lysis of remaining late stage parasites in 140 mM guanidine hydrochloride, 20mM HEPES, pH7.4 as previously described [20]. Late stage *P. knowlesi* parasites were then purified on 55% Nycodenz 72 h later. Parasites were then added to a suspension of fresh erythrocytes at ∼2 % haematocrit in complete *P. knowlesi* medium to achieve a parasitaemia of ∼2 %. Negative control cultures containing only erythrocytes were included in order to subtract background ‘parasitaemia’ counts. ML10 was then added to a final concentration of 150 nM to relevant cultures, whereas an enquivalent volume of DMSO was added to the untreated sample, and cultures incubated at 37° C for 2 h, at which time the presence of ring-stage parasites was verified by Giemsa-stained thin smears in the cultures lacking ML10. Cells were then pelleted in a microfuge at ∼2800 x g (MiniSpin^®^, Eppendorf) for 1 min before washing once with complete medium in the presence or absence of ML10 before suspending cells in complete medium for overnight incubation at 37° C in the presence or absence of ML10. Samples were set up in triplicate, with separate addition of ML10. Aliquots of the samples were fixed after the initial addition of ML10 to determine the starting parasitemia and again 17 h after washing. Fixation was performed in 0.02 % glutaraldehyde for 30 min at room temperature before storage at 4° C until required. SYBR Green I (Invitrogen) was added to aliquots of fixed parasites according to manufacturer’s instructions (10,000 x dilution) and parasites stained for 30 min at room temperature before dilution of fixed cells in 5 x volumes of PBS before parasitaemia was determined by flow cytometry using an Attune NxT.

To determine how the presence of serum in culture might affect the efficiency of ML10 to arrest egress, a synchronous culture of late-stage 3D7 parasites maintained in RPMI-1640 containing 0.5% AlbuMax was pelleted and resuspended in RPMI-1640 containing 10% human serum (without AlbuMax). The parasitemia was adjusted to 1% and the parasites were then added to wells of a 48-well plate and ML10 was added to a final concentration of 0-75 nM. Aliquots were removed and fixed to determine the starting parasitemia. The following day aliquots were removed, fixed and stained with 4.0% formaldehyde, 0.1% glutaraldehyde and SYBR Green I to determine the parasitemia of the culture by flow cytometry, as a measure of progression to the next cycle. In parallel, a culture of 3D7 that had been maintained in RPMI-1640 containing 0.5% AlbuMax and 5% human serum was treated similarly to determine the effect of human serum on the EC_50_ of ML10.

### Time-lapse video microscopy

To determine the period of time required for arrested schizonts to rupture and merozoites to egress after removal of ML10 or Compound 2, parasite cultures were maintained and assays were performed as follows. Parasites were synchronised by allowing schizonts recovered from an initial Percoll flotation to invade fresh erythrocytes for 90 minutes. The culture was subsquently treated with a second Percoll flotation and 5% D-sorbitol for 20 minutes at 37°C to retain only newly formed rings. The tightly synchronised culture of 3D7 parasites was then maintained as standard until schizonts were detected in Giemsa smears (approximately 45 hours after D-sorbitol treatment). The developed schizonts were then purified on a Percoll cushion, washed twice with medium and split equally into three different microcentrifuge tubes for a final volume of 1 ml medium in the presence of medium only (to monitor schizont development and rupture), 30 nM of ML10 or 1 µM of Compound 2 at 37°C. When late schizonts were detected and ring-stage parasites started to appear in the non-treated sample (approximately two to three hours later) the compound-arrested parasites were considered ready for washout. Meanwhile, untreated glass viewing chambers [38], medium and Lab Armor™ Beads were maintained at 37°C. When ready, parasites were spun down at 2500xg for 15 seconds, the supernatant removed and replaced with 500 µl of pre-warmed complete medium. Parasites were then added onto the pre-warmed viewing chambers and transported in a box with pre-warmed Lab Armor™ Beads. Time-lapse DIC imaging was initiated 10 minutes after addition of wash medium. Images were taken on a Nikon Ti Eclipse microscope fitted with OKO-lab environmental controls and a Hamamatsu C11440 digital camera and Nikon N Plan Apo λ 100x/1.45NA oil immersion objective. Images were taken at 5 s intervals over a total of 30 min, then exported as TIFFs or AVI, annotated and saved as mp4 files using Nikon NIS-Elements software. Videos for both ML10 and Compound 2 were in all cases taken from the same day experiment. A total of 3 independent experiments were quantified (Supp. Figure 2).

Quantification was performed using ImageJ and manually annotating in a pairwise manner the time taken for the first schizonts to egress and then the time for most schizonts in view to egress.

### Monitoring ring formation after removal of ML10 or Compound 2

To detect the formation of ring-stage parasites after release from ML10 or Compound 2 the following assay was designed. Tightly synchronised Percoll-purified schizonts were split equally into 1 mL of medium containing either 30 nM ML10 or 1 µM Compound 2 and maintained at 37°C for a maximum of 4 h, at which time the compound was removed by briefly centrifuging the cultures, aspirating the supernatant and resuspending the parasites in 1 ml of prewarmed medium at 3% haematocrit. Immediately, two 50 µl replicates were removed and fixed into 50 µl 2X fixative for a final concentration of 4% formaldehyde, 0.01% glutaraldehyde and 1X SYBR Green I and an additional 30 µl aliquot was removed for Giemsa staining; this was considered the starting time point (T=0). The remaining culture was left shaking at 37°C and samples were acquired as before after 10, 20, 30, 60 and 120 minutes. Images of Giemsa-stained smears were captured using an Olympus BX51 microscope equipped with an 100x oil objective and an Olympus SC30 camera. Comparative measurements of ring formation were obtained using an Attune NxT Flow Cytometer and analysed with Attune NxT Flow Cytometer software (ThermoFisher Scientific). SYBR Green I-positive erythrocytes were gated from a total of 200,000 singlet events and a gating strategy based on the area of SYBR Green I-positive signal was used to differentiate the schizont-stage parasites from ring-stage parasites (Supp. Figure 2). The percentage of ring-stage parasites was calculated from the total parasitaemia, i.e. all parasitized cells with a positive SYBR Green I value.

### *Survival of arrested* P. falciparum *parasites*

To determine how long *P. falciparum* parasites survive when arrested in ML10, a parasite culture was tightly synchronized using successive Percoll flotations less than two hours apart, followed by treatment with 5% D-sorbitol and subsequently incubated at 37°C. When ring-stage parasites were detected in the culture approximately 45 h later, indicating initiation of progression of the parasites to the subsequent cycle, schizont-stage parasites were isolated on a Percoll cushion. These schizonts were mixed with uninfected erythrocytes at a parasitemia of approximately 1%. ML10, Compound 2 or DMSO was immediately added to the culture to produce final concentrations of 25 nM, 1.5 μM or 0.025% (v/v) (matching the DMSO concentration of the ML10 sample), respectively. To remove the inhibitor, four 250 μl aliquots of the culture were removed, centrifuged briefly to pellet the erythrocytes. The supernatant was removed and the erythrocytes were resuspended in medium lacking inhibitor or DMSO. The resuspended parasites were subsequently placed in a well of a 48-well plate and incubated until the end of the cycle, when 50 μl of the culture was mixed with fixative solution containing 8% formaldehyde, 0.2% glutaraldehyde and 1X SYBR Green I in PBS. Fixed parasites were stored at 4°C and the parasitemia was determined using flow cytometry on an Attune NxT cytometer using Attune NxT Flow Cytometer software.

### *Growth of* P. falciparum *in the presence of ML10*

To determine the effects of ML10 on the growth of the parasites during the entire asexual erythrocytic cycle, a parasite culture was synchronized to within two hours using successive Percoll cushions and sorbitol treatment. The culture was split and either 25 nM ML10 or an equivalent volume of DMSO was then added. Synthesis of DNA was monitored by flow cyctometry using parasites that had been fixed with 4% formaldehyde and 0.1% glutaraldehyde and stained with SYBR Green I at various time points over two erythrocytic cycles. To determine if ML10-treated parasites produced invasive merozoites, ML10 and DMSO were removed 47 hours after the start of the erythrocytic cycle, three hours after the first appearance of rings in the DMSO-treated samples to ensure that the ML10 treatment covered the entire intra-erythrocytic cycle, from the earliest ring stage to the block of PKG activity.

To investigate the effect of ML10 upon continued exposure of the parasites to the compound, the growth of *P. falciparum* T618Q parasites in the presence of ML10 or DMSO over several erythrocytic cycles was measured. Cultures of *P. falciparum* T618Q and 3D7 parasites were adjusted to a parasitemia of ∼0.1%, DMSO or ML10 were added to 0.025% (v/v) and 25 nM, respectively (ensuring an equal concentration of DMSO in each sample) and growth was subsequently measured over four cycles. Starting four days after the addition of the compound, the medium was replaced with fresh medium containing DMSO or ML10 daily. The parasitemia was measured by flow cytometry as described above.

### Statistical analysis

Statiscal analyses were performed using GraphPad Prism 8 software and significance is represented as *** *p*≤0.001, \*\**p*≤0.01 and \**p*≤0.05. The different statistical analyses applied in this study are described in the respective figure legends, when applicable.

## Acknowledgments

We thank Avnish Patel for constructive feedback on experimental design, Donelly van Schalkwyk for providing Dd2 parasites, Julian Muwanguzi-Karugaba for providing HB3 parasites and Michael Delves for providing human serum. This work was supported by Medical Reseach Council Career Development Awards MR/R008485/1 (CvO) and MR/M021157/1 (RWM) (jointly funded by the UK MRC and the UK Department for International Development), Wellcome Trust grant 106240/Z/14/Z (DAB), British Biological Sciences Research Council grant BB/M009513/1 (SDN) and Wellcome Trust Sir Henry Wellcome fellowship 210861/Z/18/Z (JAT).

## FIGURE LEGENDS

**Supplementary Figure 1. Scatter plots of the mean EC**_**50**_ **values obtained from multiple independent dose-response curves with different *P. falciparum* lines and culture conditions and *P. knowlesi***. Mean EC_50_ values for *P. falciparum* strains 3D7, Dd2, HB3 and NF54 grown in standard culture conditions (0.5% AlbuMax) in the presence of ML10 (**A**) or Compound 2 (**B**). (**C**) Mean EC_50_ values obtained with *P. falciparum* strain 3D7 grown in standard culture conditions (0.5% AlbuMax) or in RPMI-1640 containing 0.5% AlbuMax and 10% horse serum. (**D**) Mean EC_50_ values obtained with *P. knowlesi* strain A1-H.1 grown in standard culture conditions using RPMI-1640 containing 0.5% AlbuMax and 10% horse serum. Error bars are standard deviation of the calculated means.

**Supplementary Figure 2. Pairwise scatter plots depicting the start and duration of egress after removal of ML10 or Compound 2** Quantification of the period until the first schizont egress event was detected (**A**) and time required for egress of the majority of schizonts after egression of the first schizont (**B**). Values were attained from the analysis of egress videos after removal of ML10 or Compound 2 from 3 independent experiments. No significant differences were detected using a paired t-test.

**Supplementary Figure 3. Gating strategy for determination of rings after compound washout represented in Figure 5**. Representative gating strategies for the determination of parasitized erythrocytes stained with SYBR Green I nuclear staining and their respective ring and schizont stage, 10 minutes (**A**) and 60 minutes after (**B**) compound washout. (**C**) Overlay of histograms of the two time points emphasizing the differences in signal profiles between ring and schizont populations.

**Supplementary Video 1. Video microscopy of egress after removal of ML10** Egress of *P. falciparum* merozoites *in vitro* imaged by time-lapse video microscopy following removal of ML10.

**Supplementary Video 2. Video microscopy of egress after removal of Compound 2** Egress of *P. falciparum* merozoites *in vitro* imaged by time-lapse video microscopy following removal of Compound 2.

## REFERENCES

1. Bozdech Z, Llinas M, Pulliam BL, Wong ED, Zhu J, DeRisi JL. The transcriptome of the intraerythrocytic developmental cycle of *Plasmodium falciparum*. PLoS Biol. 2003;1(1):E5.

2. Le Roch KG, Zhou Y, Blair PL, Grainger M, Moch JK, Haynes JD, et al. Discovery of gene function by expression profiling of the malaria parasite life cycle. Science. 2003;301(5639):1503–8.

3. Llinas M, Bozdech Z, Wong ED, Adai AT, DeRisi JL. Comparative whole genome transcriptome analysis of three *Plasmodium falciparum* strains. Nucleic Acids Res. 2006;34(4):1166–73.

4. Chua ACY, Ong JJY, Malleret B, Suwanarusk R, Kosaisavee V, Zeeman AM, et al. Robust continuous *in vitro* culture of the *Plasmodium cynomolgi* erythrocytic stages. Nat Commun. 2019;10(1):3635.

5. Lim C, Hansen E, DeSimone TM, Moreno Y, Junker K, Bei A, et al. Expansion of host cellular niche can drive adaptation of a zoonotic malaria parasite to humans. Nat Commun. 2013;4:1638.

6. Moon RW, Hall J, Rangkuti F, Ho YS, Almond N, Mitchell GH, et al. Adaptation of the genetically tractable malaria pathogen *Plasmodium knowlesi* to continuous culture in human erythrocytes. Proc Natl Acad Sci U S A. 2013;110(2):531–6.

7. Trager W, Jensen JB. Human malaria parasites in continuous culture. Science. 1976;193(4254):673–5.

8. Hirako IC, Assis PA, Hojo-Souza NS, Reed G, Nakaya H, Golenbock DT, et al. Daily Rhythms of TNFalpha Expression and Food Intake Regulate Synchrony of *Plasmodium* Stages with the Host Circadian Cycle. Cell Host Microbe. 2018;23(6):796–808 e6.

9. Mideo N, Reece SE, Smith AL, Metcalf CJ. The Cinderella syndrome: why do malaria-infected cells burst at midnight? Trends Parasitol. 2013;29(1):10–6.

10. Kwiatkowski D, Greenwood BM. Why is malaria fever periodic? A hypothesis. Parasitol Today. 1989;5(8):264–6.

11. O’Donnell AJ, Schneider P, McWatters HG, Reece SE. Fitness costs of disrupting circadian rhythms in malaria parasites. Proc Biol Sci. 2011;278(1717):2429–36.

12. Glushakova S, Beck JR, Garten M, Busse BL, Nasamu AS, Tenkova-Heuser T, et al. Rounding precedes rupture and breakdown of vacuolar membranes minutes before malaria parasite egress from erythrocytes. Cell Microbiol. 2018;20(10):e12868.

13. Hale VL, Watermeyer JM, Hackett F, Vizcay-Barrena G, van Ooij C, Thomas JA, et al. Parasitophorous vacuole poration precedes its rupture and rapid host erythrocyte cytoskeleton collapse in *Plasmodium falciparum* egress. Proc Natl Acad Sci U S A. 2017;114(13):3439–44.

14. Thomas JA, Tan MSY, Bisson C, Borg A, Umrekar TR, Hackett F, et al. A protease cascade regulates release of the human malaria parasite *Plasmodium falciparum* from host red blood cells. Nat Microbiol. 2018;3(4):447–55.

15. Weiss GE, Gilson PR, Taechalertpaisarn T, Tham WH, de Jong NW, Harvey KL, et al. Revealing the sequence and resulting cellular morphology of receptor-ligand interactions during *Plasmodium falciparum* invasion of erythrocytes. PLoS Pathog. 2015;11(2):e1004670.

16. Inselburg J, Banyal HS. Plasmodium falciparum: synchronization of asexual development with aphidicolin, a DNA synthesis inhibitor. Exp Parasitol. 1984;57(1):48–54.

17. Assaraf YG, Golenser J, Spira DT, Bachrach U. *Plasmodium falciparum*: synchronization of cultures with DL-alpha-difluoromethylornithine, an inhibitor of polyamine biosynthesis. Exp Parasitol. 1986;61(2):229–35.

18. van Biljon R, Niemand J, van Wyk R, Clark K, Verlinden B, Abrie C, et al. Inducing controlled cell cycle arrest and re-entry during asexual proliferation of *Plasmodium falciparum* malaria parasites. Sci Rep. 2018;8(1):16581.

19. Lambros C, Vanderberg JP. Synchronization of *Plasmodium falciparum* erythrocytic stages in culture. J Parasitol. 1979;65(3):418–20.

20. Ngernna S, Chim-Ong A, Roobsoong W, Sattabongkot J, Cui L, Nguitragool W. Efficient synchronization of *Plasmodium knowlesi in vitro* cultures using guanidine hydrochloride. Malar J. 2019;18(1):148.

21. Brown AC, Moore CC, Guler JL. Cholesterol-dependent enrichment of understudied erythrocytic stages of human *Plasmodium parasites*. Sci Rep. 2020;10(1):4591.

22. Geislinger TM, Chan S, Moll K, Wixforth A, Wahlgren M, Franke T. Label-free microfluidic enrichment of ring-stage *Plasmodium falciparum*-infected red blood cells using non-inertial hydrodynamic lift. Malar J. 2014;13:375.

23. Tosta CE, Sedegah M, Henderson DC, Wedderburn N. *Plasmodium yoelii* and *Plasmodium berghei*: isolation of infected erythrocytes from blood by colloidal silica gradient centrifugation. Exp Parasitol. 1980;50(1):7–15.

24. Saul A, Myler P, Elliott T, Kidson C. Purification of mature schizonts of *Plasmodium falciparum* on colloidal silica gradients. Bull World Health Organ. 1982;60(5):755–9.

25. Rivadeneira EM, Wasserman M, Espinal CT. Separation and concentration of schizonts of *Plasmodium falciparum* by Percoll gradients. J Protozool. 1983;30(2):367–70.

26. Kramer KJ, Kan SC, Siddiqui WA. Concentration of *Plasmodium falciparum*-infected erythrocytes by density gradient centrifugation in Percoll. J Parasitol. 1982;68(2):336–7.

27. Radfar A, Mendez D, Moneriz C, Linares M, Marin-Garcia P, Puyet A, et al. Synchronous culture of *Plasmodium falciparum* at high parasitemia levels. Nat Protoc. 2009;4(12):1899–915.

28. Childs RA, Miao J, Gowda C, Cui L. An alternative protocol for *Plasmodium falciparum* culture synchronization and a new method for synchrony confirmation. Malar J. 2013;12:386.

29. Mata-Cantero L, Lafuente MJ, Sanz L, Rodriguez MS. Magnetic isolation of *Plasmodium falciparum* schizonts iRBCs to generate a high parasitaemia and synchronized *in vitro* culture. Malar J. 2014;13:112.

30. Trang DT, Huy NT, Kariu T, Tajima K, Kamei K. One-step concentration of malarial parasite-infected red blood cells and removal of contaminating white blood cells. Malar J. 2004;3:7.

31. Paul F, Roath S, Melville D, Warhurst DC, Osisanya JO. Separation of malaria-infected erythrocytes from whole blood: use of a selective high-gradient magnetic separation technique. Lancet. 1981;2(8237):70–1.

32. Ranford-Cartwright LC, Sinha A, Humphreys GS, Mwangi JM. New synchronization method for *Plasmodium falciparum*. Malar J. 2010;9:170.

33. Boyle MJ, Wilson DW, Richards JS, Riglar DT, Tetteh KK, Conway DJ, et al. Isolation of viable *Plasmodium falciparum* merozoites to define erythrocyte invasion events and advance vaccine and drug development. Proc Natl Acad Sci U S A. 2010;107(32):14378-83. Epub 2010/07/28.

34. Lyth O, Vizcay-Barrena G, Wright KE, Haase S, Mohring F, Najer A, et al. Cellular dissection of malaria parasite invasion of human erythrocytes using viable *Plasmodium knowlesi* merozoites. Sci Rep. 2018;8(1):10165.

35. Dennis ED, Mitchell GH, Butcher GA, Cohen S. *In vitro* isolation of *Plasmodium knowlesi* merozoites using polycarbonate sieves. Parasitology. 1975;71(3):475–81.

36. Knuepfer E, Suleyman O, Dluzewski AR, Straschil U, O’Keeffe AH, Ogun SA, et al. RON12, a novel *Plasmodium*-specific rhoptry neck protein important for parasite proliferation. Cell Microbiol. 2014;16(5):657–72.

37. Alam MM, Solyakov L, Bottrill AR, Flueck C, Siddiqui FA, Singh S, et al. Phosphoproteomics reveals malaria parasite Protein Kinase G as a signalling hub regulating egress and invasion. Nat Commun. 2015;6:7285.

38. Collins CR, Hackett F, Strath M, Penzo M, Withers-Martinez C, Baker DA, et al. Malaria parasite cGMP-dependent protein kinase regulates blood stage merozoite secretory organelle discharge and egress. PLoS Pathog. 2013;9(5):e1003344.

39. Diaz CA, Allocco J, Powles MA, Yeung L, Donald RG, Anderson JW, et al. Characterization of *Plasmodium falciparum* cGMP-dependent protein kinase (PfPKG): antiparasitic activity of a PKG inhibitor. Mol Biochem Parasitol. 2006;146(1):78–88.

40. Taylor HM, McRobert L, Grainger M, Sicard A, Dluzewski AR, Hopp CS, et al. The malaria parasite cyclic GMP-dependent protein kinase plays a central role in blood-stage schizogony. Eukaryot Cell. 2010;9(1):37–45.

41. Donald RG, Zhong T, Wiersma H, Nare B, Yao D, Lee A, et al. Anticoccidial kinase inhibitors: identification of protein kinase targets secondary to cGMP-dependent protein kinase. Mol Biochem Parasitol. 2006;149(1):86–98.

42. Collins CR, Hackett F, Atid J, Tan MSY, Blackman MJ. The *Plasmodium falciparum* pseudoprotease SERA5 regulates the kinetics and efficiency of malaria parasite egress from host erythrocytes. PLoS Pathog. 2017;13(7):e1006453.

43. Sherling ES, Knuepfer E, Brzostowski JA, Miller LH, Blackman MJ, van Ooij C. The *Plasmodium falciparum* rhoptry protein RhopH3 plays essential roles in host cell invasion and nutrient uptake. Elife. 2017;6.

44. Mohring F, Hart MN, Rawlinson TA, Henrici R, Charleston JA, Diez Benavente E, et al. Rapid and iterative genome editing in the malaria parasite *Plasmodium knowlesi* provides new tools for *P. vivax* research. Elife. 2019;8.

45. Pino P, Caldelari R, Mukherjee B, Vahokoski J, Klages N, Maco B, et al. A multistage antimalarial targets the plasmepsins IX and X essential for invasion and egress. Science. 2017;358(6362):522–8.

46. Baker DA, Stewart LB, Large JM, Bowyer PW, Ansell KH, Jimenez-Diaz MB, et al. A potent series targeting the malarial cGMP-dependent protein kinase clears infection and blocks transmission. Nat Commun. 2017;8(1):430.

47. Smilkstein M, Sriwilaijaroen N, Kelly JX, Wilairat P, Riscoe M. Simple and inexpensive fluorescence-based technique for high-throughput antimalarial drug screening. Antimicrob Agents Chemother. 2004;48(5):1803–6.

48. Gurnett AM, Liberator PA, Dulski PM, Salowe SP, Donald RG, Anderson JW, et al. Purification and molecular characterization of cGMP-dependent protein kinase from Apicomplexan parasites. A novel chemotherapeutic target. J Biol Chem. 2002;277(18):15913–22.

49. Chugh M, Scheurer C, Sax S, Bilsland E, van Schalkwyk DA, Wicht KJ, et al. Identification and deconvolution of cross-resistance signals from antimalarial compounds using multidrug-resistant *Plasmodium falciparum* strains. Antimicrob Agents Chemother. 2015;59(2):1110–8.

50. Murray L, Stewart LB, Tarr SJ, Ahouidi AD, Diakite M, Amambua-Ngwa A, et al. Multiplication rate variation in the human malaria parasite *Plasmodium falciparum*. Sci Rep. 2017;7(1):6436.

51. van Schalkwyk DA, Moon RW, Blasco B, Sutherland CJ. Comparison of the susceptibility of *Plasmodium knowlesi* and *Plasmodium falciparum* to antimalarial agents. J Antimicrob Chemother. 2017;72(11):3051–8.

52. van Schalkwyk DA, Blasco B, Davina Nunez R, Liew JWK, Amir A, Lau YL, et al. *Plasmodium knowlesi* exhibits distinct in vitro drug susceptibility profiles from those of *Plasmodium falciparum*. Int J Parasitol Drugs Drug Resist. 2019;9:93–9.

53. Boyle MJ, Richards JS, Gilson PR, Chai W, Beeson JG. Interactions with heparin-like molecules during erythrocyte invasion by *Plasmodium falciparum* merozoites. Blood. 2010;115(22):4559–68.

54. Das S, Hertrich N, Perrin AJ, Withers-Martinez C, Collins CR, Jones ML, et al. Processing of *Plasmodium falciparum* Merozoite Surface Protein MSP1 Activates a Spectrin-Binding Function Enabling Parasite Egress from RBCs. Cell Host Microbe. 2015;18(4):433-44. Epub 2015/10/16.

55. Patel A, Perrin AJ, Flynn HR, Bisson C, Withers-Martinez C, Treeck M, et al. Cyclic AMP signalling controls key components of malaria parasite host cell invasion machinery. PLoS Biol. 2019;17(5):e3000264.

56. Flueck C, Drought LG, Jones A, Patel A, Perrin AJ, Walker EM, et al. Phosphodiesterase beta is the master regulator of cAMP signalling during malaria parasite invasion. PLoS Biol. 2019;17(2):e3000154. Epub 2019/02/23.

57. Bannister LH, Hopkins JM, Margos G, Dluzewski AR, Mitchell GH. Three-dimensional ultrastructure of the ring stage of *Plasmodium falciparum*: evidence for export pathways. Microsc Microanal. 2004;10(5):551–62.

58. Gruring C, Heiber A, Kruse F, Ungefehr J, Gilberger TW, Spielmann T. Development and host cell modifications of *Plasmodium falciparum* blood stages in four dimensions. Nat Commun. 2011;2:165.

59. Collins CR, Das S, Wong EH, Andenmatten N, Stallmach R, Hackett F, et al. Robust inducible Cre recombinase activity in the human malaria parasite *Plasmodium falciparum* enables efficient gene deletion within a single asexual erythrocytic growth cycle. Mol Microbiol. 2013;88(4):687–701.

60. Portugaliza HP, Llora-Batlle O, Rosanas-Urgell A, Cortes A. Reporter lines based on the *gexp02* promoter enable early quantification of sexual conversion rates in the malaria parasite *Plasmodium falciparum*. Sci Rep. 2019;9(1):14595.

61. Perrin AJ, Collins CR, Russell MRG, Collinson LM, Baker DA, Blackman MJ. The Actinomyosin Motor Drives Malaria Parasite Red Blood Cell Invasion but Not Egress. MBio. 2018;9(4).

62. Chang CY, Simon E. The effect of dimethyl sulfoxide (DMSO) on cellular systems. Proc Soc Exp Biol Med. 1968;128(1):60–6.

63. Lampugnani MG, Pedenovi M, Niewiarowski A, Casali B, Donati MB, Corbascio GC, et al. Effects of dimethyl sulfoxide (DMSO) on microfilament organization, cellular adhesion, and growth of cultured mouse B16 melanoma cells. Exp Cell Res. 1987;172(2):385–96.

64. Moon RW, Sharaf H, Hastings CH, Ho YS, Nair MB, Rchiad Z, et al. Normocyte-binding protein required for human erythrocyte invasion by the zoonotic malaria parasite *Plasmodium knowlesi*. Proc Natl Acad Sci U S A. 2016;113(26):7231–6.

