## Supplementary figures and images for "Use of a highly specific kinase inhibitor for rapid, simple and precise synchronization of *Plasmodium falciparum* and *Plasmodium knowlesi* asexual stage parasites"

### Supplemental Data 1

#### Supp. figure 1

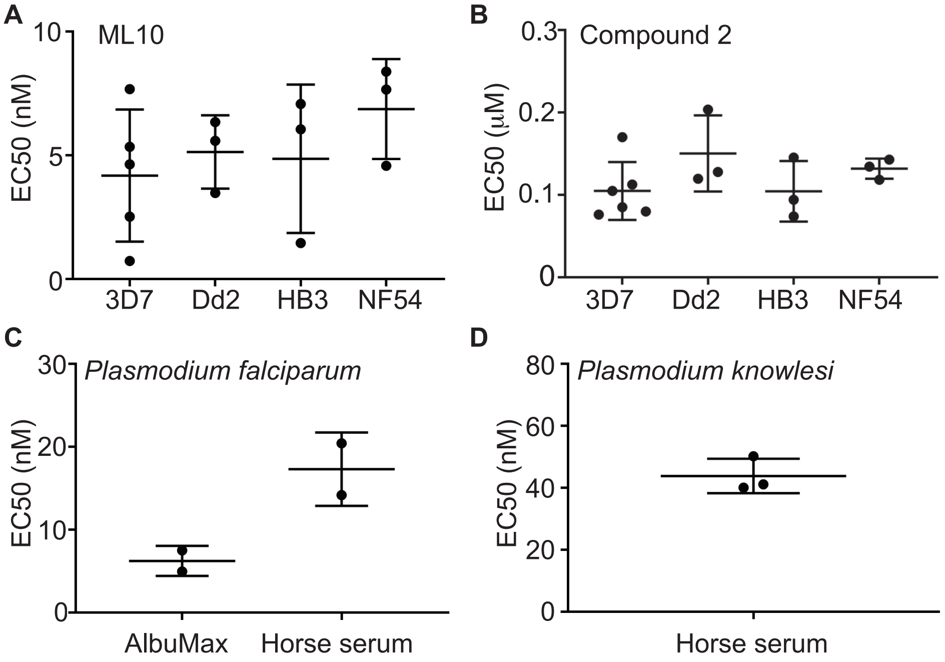

#### Supp. figure 2

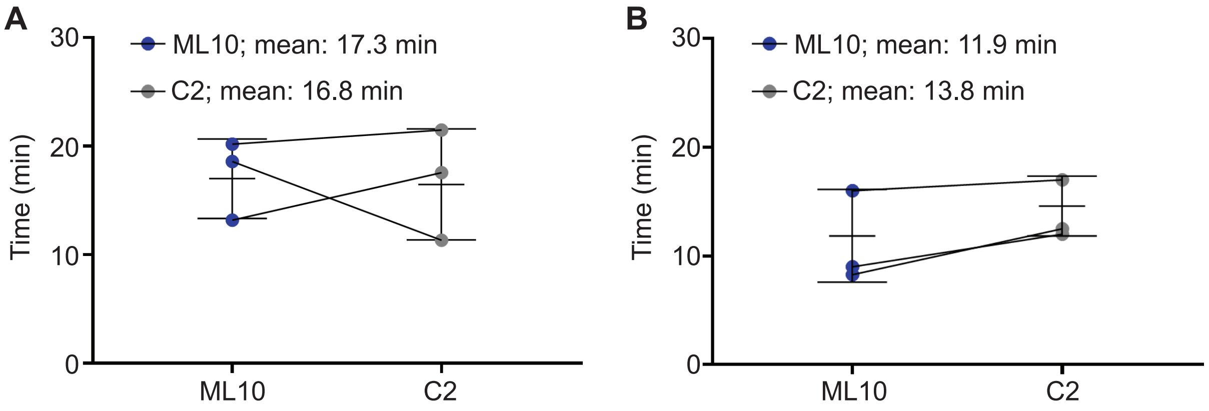


#### Supp. figure 3

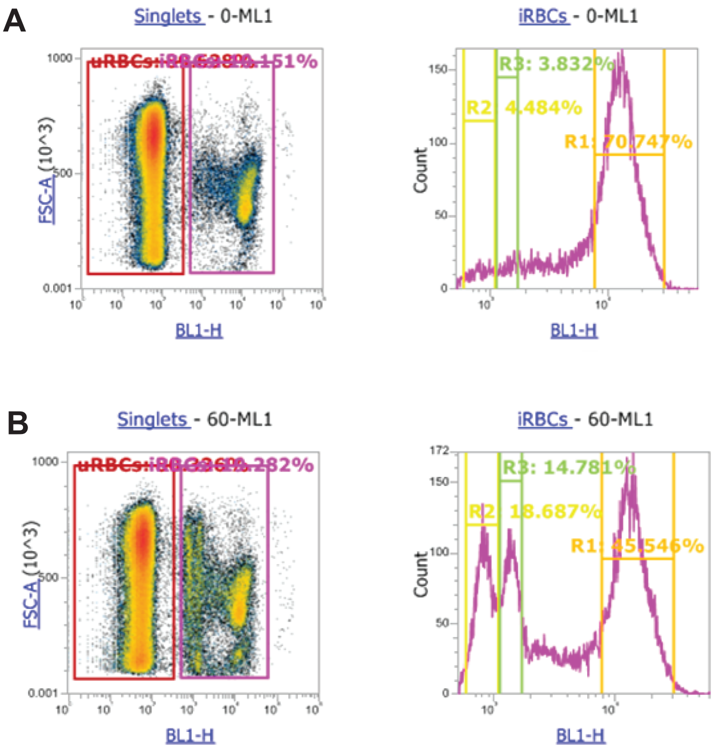
